# Gliding motility of a uranium tolerant *Bacteroidetes* bacterium *Chryseobacterium* sp. strain PMSZPI: Insights into the architecture of spreading colonies

**DOI:** 10.1101/2021.07.27.453926

**Authors:** Devanshi Khare, Pallavi Chandwadkar, Celin Acharya

## Abstract

Uranium tolerant soil bacterium *Chryseobacterium* sp. strain PMSZPI moved over solid agar surfaces by gliding motility thereby forming spreading colonies which is a hallmark of members of Bacteroidetes phylum. PMSZPI genome harbored orthologs of all the *gld* and *spr* genes considered as core bacteroidetes gliding motility genes of which *gldK, gldL, gldM,* and *gldN* were co-transcribed. Here, we present the intriguing interplay between gliding motility and cellular organization in PMSZPI spreading colonies. While nutrient deficiency enhanced colony spreading, high agar concentrations and presence of motility inhibitor like 5-hydroxyindole reduced the spreading. A detailed *in situ* structural analysis of spreading colonies revealed closely packed cells forming multiple layers at center of colony while the edges showed clusters of cells periodically arranged in hexagonal lattices interconnected with each other. The cell migration within the colony was visualized as branched structures wherein the cells were buried within extracellular matrix giving rise to ‘fern’ like patterns. PMSZPI colonies exhibited strong iridescence that showed correlation with gliding motility. Presence of uranium reduced motility and iridescence and induced biofilm formation. This is a first report of gliding motility and iridescence in a bacterium from uranium enriched environment that could be of significant interest from an ecological perspective.

**Originality-Significance Statement:** This work provides the first description of the gliding motility and iridescence or structural coloration in a Bacteroidetes soil bacterium from uranium enriched environment. The periodic arrangement of the cell population in the spreading colonies achieved through gliding motility resulted in bright structural coloration of the colonies when illuminated. The study describes the exogenous factors including nutrition, substrate, presence of uranium influencing the motility and iridescence of the bacterium. The highly organized cell population in the gliding and iridescent bacterium may have conferred survival advantage in metal/uranium enriched ecosystem.

## Introduction

Bacteria compete with each other for resources and space and employ ingenious mechanisms to successfully occupy and establish their niche. Bacterial motility is a universal phenotypic attribute that allows various lifestyles and ecological adaptation. Motility allows the bacteria to escape stresses or facilitates movement toward nutrients ensuring their survival (Wei *et al*., 2011). Surfaces form one of the most important territories of microbial life (Kolter and Greenberg, 2006) and the microbial surface motility allows some species to rapidly colonize surfaces initiating biofilm formation (Dang and Lovell, 2016). Swarming, gliding, twitching or sliding modes of bacterial surface translocation offer advantages in survival and competition (O’Toole and Kolter, 1998; Jarrell and McBride, 2008; Kearns, 2010)

The phylum Bacteroidetes comprises of a wide variety of Gram-negative, rod shaped bacteria that inhabit several ecosystems ranging from aquatic, soil, sediment, terrestrial to the gut microflora (Hahnke *et al*., 2016). The members of Bacteroidetes are known to navigate surfaces by a unique form of motility, known as gliding motility, which occurs without the aid of any external organelle like pili and flagella (Jarrell and McBride, 2008). Gliding motility enables the movement of the bacteria along the solid surfaces and results in spreading colonies (Penttinen *et al*., 2018). Gliding motility in Bacteroidetes has largely been studied in Cytophagales and the Flavobacteriales contributing towards nutrient acquisition and colonization (McBride, 2001; Kita *et al*., 2016). Some proteins required for gliding are components of a novel protein secretion system, the Type IX Secretion System (T9SS) or the Por Secretion System (Sato *et al*., 2010; McBride and Zhu, 2013). *Flavobacterium johnsoniae,* a non-pathogenic strain, that is commonly found in freshwater and soil has emerged as a robust model system for studying the mechanism of gliding motility specific to Bacteroidetes. Molecular analyses identified 19 genes involved in *F. johnsoniae* gliding motility-the *gld* genes (*gldA*, *gldB, gldD, gldF, gldG, gld H, gldI, gldJ, gldK, gldL, gldM, gldN*) that are essential for gliding and the *spr* genes (*sprA, sprB, sprC, sprD, sprE, sprF, sprT*) that are important but not entirely essential for gliding (Agarwal *et al*., 1997; Hunnicutt and McBride, 2000, 2001; McBride and Zhu, 2013; McBride and Nakane, 2015). Furthermore, a subset of these genes, *gldK, gldL, gldM, gldN, sprA, sprE,* and *sprT* constitutes the T9SS (Sato *et al*., 2010; McBride and Zhu, 2013) which is specific to Bacteroidetes with no similarity with the previously defined bacterial secretion systems ranging from Type I to Type VI and Type VIII (McBride and Zhu, 2013). Gliding motility was shown to contribute towards the maintenance of the periodicity within the cell population of biofilms with iridescent properties in *Cellulophaga* spp. (Kientz *et al*., 2016).

We recently studied the genomic and functional attributes of a uranium tolerant Bacteroidetes bacterium, *Chryseobacterium* sp. strain PMSZPI (Khare *et al*., 2020) that was isolated from the sub-surface soil of a uranium ore deposit (Kumar *et al*., 2013). The genus *Chryseobacterium* belonging to the family *Flavobacteriaceae* was separated from the genus *Flavobacterium* to provide it a distinct taxonomic status (Vandamme *et al*., 1994; Bernardet *et al*., 1996). PMSZPI demonstrated a wide range of adaptation and resistance strategies which apparently allowed its survival enduring an ecological system comprising of high concentrations of uranium and other heavy metals. The strain was shown to be motile via gliding motility (Khare *et al*., 2020). In this study, we present the characteristics of gliding motility under various growth and substrate conditions at colonial level. The structural organization of the cells in the spreading colony was analyzed in detail in order to gain insights into the features contributing to the optical appearance of the colony. Our studies present the intriguing interplay among the gliding motility, cellular organization and iridescence in this uranium tolerant bacterium. Furthermore, implications of uranium exposure on the gliding motility, biofilm formation and iridescence were also explored and the results suggested that the presence of uranium is an important regulator of both gliding motility and iridescence.

## Results and discussion

### *Chryseobacterium* PMSZPI encodes core Bacteroidetes gliding motility genes

*Chryseobacterium* sp. strain PMSZPI is a Gram-negative, metal tolerant, rod-shaped bacterium belonging to the phylum Bacteroidetes that demonstrated gliding motility on agar surfaces (Khare *et al*., 2020). Although gliding motility has been reported amongst Bacteroidetes members, extensive studies on mechanisms of gliding motility are limited to *Flavobacterium johnsoniae. Chryseobacterium* PMSZPI was found to be distantly related to *F. johnsoniae* (Fig. 1A). Transmission electron microscopic analysis could not identify any motility machines like pili and flagella on the cells of PMSZPI (Fig. 1B) which are known to facilitate the movement of cells over surfaces in other bacterial strains (Harshey, 1994; Mattick, 2002). Moreover, the genome analysis also failed to categorize genes encoding the essential components of flagella and type IV pili in *Chryseobacterium* PMSZPI. Therefore, it was anticipated that the gliding motility in PMSZPI relied on motility machinery other than flagella or pili.

**Figure 1:**
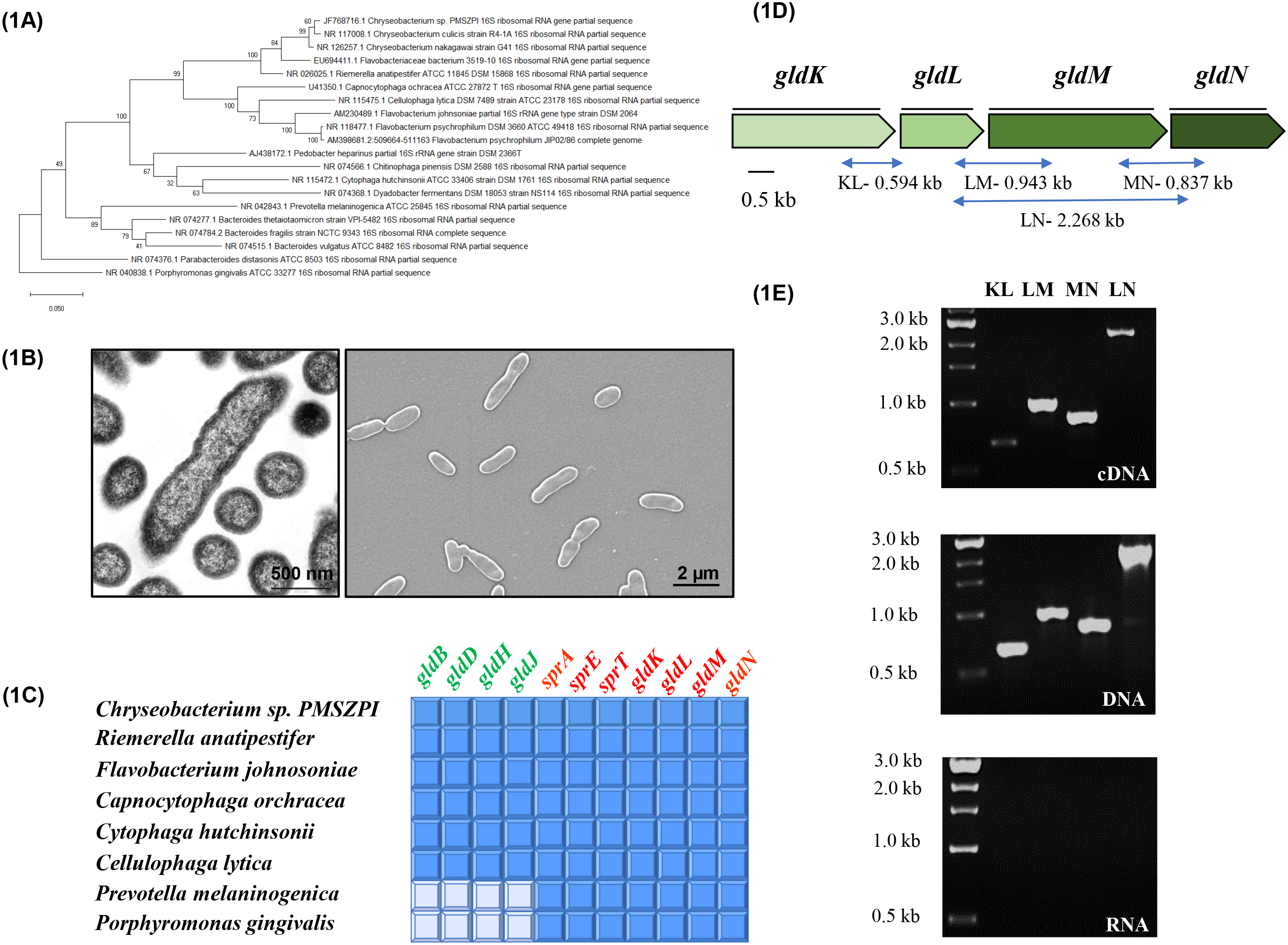
Phylogenetic affiliation of *Chryseobacterium* sp. strain PMSZPI and its gliding motility genes. **(A)** Maximum Likelihood phylogram of phylum Bacteroidetes based on 16S rRNA gene sequence data. Phylogenetic tree was generated using MEGA 7 package with 500 bootstrap replications. (B) Transmission electron micrograph and Scanning electron micrograph displaying the rod-shaped cells of PMSZPI without any appendages. (C) Orthologs to the core Bacteroidetes gliding motility genes of *F. johnsoniae* in the members of Bacteroidetes. The orthologs were identified by BLAST analysis and their presence in the various genomes are depicted by coloured box and absence by white box. (D) Schematic representation of arrangement of *gldK, gldL, gldM* and *gldN* genes in PMSZPI genome. Regions of the gene and their sizes (in kb) which have been amplified in Fig. 1E are mentioned. (E) RT-PCR of *Chryseobacterium* sp. PMSZPI RNA. Reverse transcription was performed with primers covering internal regions of two adjacent genes (KL, LM, MN) and LN. For each region amplified, three reactions were electrophoresed on 1% agarose gel; RT-PCR mixture with cDNA as template (gel 1), positive-control PCR mixture with PMSZPI genomic DNA as template (gel 2) and negative control PCR mixture with RNA as template (gel 3). First lane of each gel represents 1 kb DNA ladder (NEB), Lanes 2, 3, 4 and 5 correspond to regions amplified by primers for internal regions of KL, LM, MN and LN respectively for all the gels.

Several related members of *F. johnsoniae* within Bacteroidetes have orthologs for the *gld* and *spr* genes and show rapid gliding motility. Analysis of PMSZPI genome revealed orthologs of fifteen genes (*gldA, gldB, gldF, gldD, gldH, gldI, gldE, gldJ, gldK, gldL, gldM, gldN, sprA, sprE, sprT*) that are reported to be involved in gliding motility in *F. johnsoniae* (Sato *et al*., 2010; McBride and Nakane, 2015). The cell envelope proteins GldK, GldL, GldM, GldN, SprA, SprE and SprT are central components of the type IX secretion system (T9SS) which are crucial for gliding motility machinery in *F. johnsoniae* (McBride and Zhu, 2013). Orthologs to eleven genes categorized as core bacteroidetes gliding motility genes - *gldB, gldD, gldH, gldJ, gldK, gldL, gldM, gldN, sprA, sprE* and *sprT* (McBride and Zhu, 2013) were identified in PMSZPI alongside with other gliding members of *Bacteroidetes* except *Porphyromonas gingivalis* and *Prevotella melaninogenica* which lacked gliding motility genes and are non-motile (Fig. 1C, Table S1A). Originally, *gldK, gldL, gldM* and *gldN* genes were discovered as gliding motility genes of *F. johnsoniae* (Braun *et al*., 2005). Orthologs of these genes are present in many Bacteroidetes members which seem to be consistently clustered together on the genomes indicating the co-transcription of these genes (Shrivastava *et al*., 2013). The sizes of *gldK, gldL, gldM and gldN* of PMSZPI were comparable to that of other Bacteroidetes strains (Table S1B). Phylogenetic relationship analysis of GldK, GldL, GldM, and GldN sequences using maximum likelihood method revealed the closest relationship between PMSZPI and *Riemerella anatipestifer* for the orthologous proteins (Fig. S1). The profile of phylogenetic trees for GldK, GldL, GldM, and GldN sequences were similar to those based on 16S rRNAs (Figs. 1A and S1) indicating that these genes were most likely transferred vertically. In order to evaluate the transcriptional organization of *gldK, gldL, gldM and gldN* in PMSZPI, reverse transcriptase PCR (RT-PCR) was employed using oligonucleotides (Table S2) in various combinations namely KL, LM, MN and LN respectively for the amplification of internal regions (Fig. 1D) and total purified RNA extracted from PMSZPI cells. RT-PCR products with the expected sizes were obtained for each gene junction of the *gldKLMN* gene cluster from DNA or cDNA suggesting that these four genes were co-transcribed (Fig. 1E). The genetic organization of *gldK, gldL, gldM,* and *gldN* amongst Bacteroidetes members is said to be conserved for their function as their coordinated expression is apparently required for efficient assembly of T9SS complex (Shrivastava *et al*., 2013).

### Colony spreading of *Chryseobacterium* PMSZPI is influenced by incubation time and concentrations of nutrient, agar and motility inhibitor

Bacteroidetes cells exhibiting gliding motility typically form spreading colonies (McBride, 2001; McBride and Nakane, 2015) which is a vital phenotypic indicator of the intact and active gliding motility system. The morphology of macrocolonies of PMSZPI was assayed on LB medium containing 0.35% agar as previously shown for illustrating gliding motility in Bacteroidetes members (Li *et al*., 2015). The exponential phase PMSZPI cells (10 µL) were spotted on soft agar (0.35%), incubated at 30°C and the colony morphology was imaged at various time intervals. The cells spread radially from all the directions of the inoculated site forming large colonies that showed irregular spreading edges (Fig 2A). A progressive colony expansion (∼89%) was visualized over a period of seven days (Fig. 2A) indicating that the development of the colonies depended on incubation time. Motility is important in organisms which apparently allows them to move towards the optimal concentrations of nutrients. To evaluate the effect of nutrient concentration on gliding motility, we compared the gliding behaviour of PMSZPI cells on soft agar (0.35%) with 1/2 to 1/50 strength of LB medium concentration. The size of the developing colonies increased on nutrient-deficient medium with maximum colony spreading observed on 1/50 LB medium over 24 h of incubation (Fig. 2B) suggesting that the gliding motility performance was enhanced by nutrient deprivation. Highly intricate dendritic branching patterns in the colonies emanating from site of inoculation were observed with lower strength of LB concentrations (1/10-1/50) (Fig. 2B) in contrast to normal LB concentration wherein no such branching patterns were visualized (Fig. 2A) possibly due to higher cell densities on nutrient rich medium. Nutrient-poor conditions have been shown to favor motility and colony spreading in Flavobacteria (Harshey, 1994; Penttinen *et al*., 2018). The correlation of the gliding performance with the physical strength of the culture substrate was observed by inoculating PMSZPI cells on to LB medium (1/10 LB was taken for optimal spreading) with agar concentrations varying from 0.35% to 1%. Colony spreading decreased as the agar concentration increased in the medium with PMSZPI forming circular colonies on 0.7% and 1% agar concentrations that hardly showed any spreading beyond the inoculation spots. It could be for the reason that the cell motility reduced with increase in agar concentration suggesting that the motility was higher on soft substrate (Fig. 2C). This was in contrast to gliding motility phenotype of *F. johnsoniae* wherein the motility was low on the soft agar substrate (Sato *et al*., 2021). The colony spreading or the motility of PMSZPI was indistinguishable in the presence or absence of glucose (0.2-0.4%) to 0.35% LB agar medium (data not included). Under all the conditions studied here, colony spreading of PMSZPI appeared in two stages-an early growth-dependent phase followed by a secondary gliding motility dependent phase which caused spreading and colony expansion. This kind of colony spreading in PMSZPI was similar to that of *F. johnsoniae (Sato et al., 2021)*.

**Figure 2:**
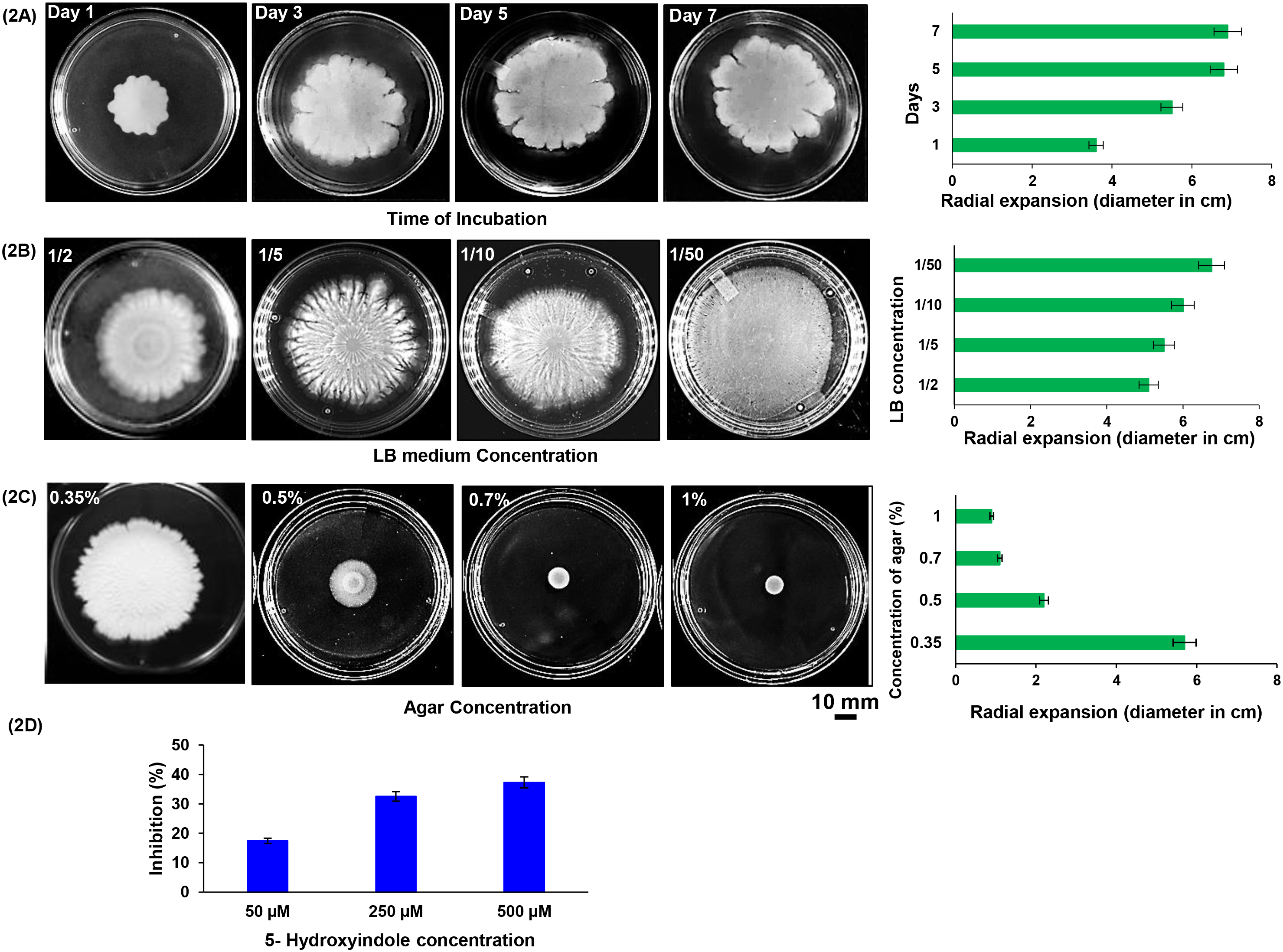
Colony spreading of PMSZPI at different incubation periods and concentrations of nutrient medium, agar and motility inhibitor. PMSZPI cells (10 µl at cell density of 2 x 10^5^ total cells) were spotted on (A) LB 0.35% agar and incubated at 30°C until 7 d to determine the effect of incubation time on colony spreading. Images of day wise motility in petri plates and the corresponding histogram showing the change in colony diameters with increase in incubation time are shown. (B) LB medium ranging from 1/2, 1/5, 1/10 and 1/50 strength concentrations with 0.35% agar to determine the effect of nutrient concentrations on colony spreading and incubated at 30°C for 1 d. The corresponding histogram showing increase in diameter with decrease in LB concentrations is shown. (C) LB (1/10) containing 0.35-1% agar and incubated at 30°C for 1 d to study the effect of agar concentrations on colony spreading. The corresponding histogram depicting the decrease in colony diameter with increase in agar concentrations is shown. (D) LB 0.35% agar supplemented with motility inhibitor, 5-Hydroxyindole at concentrations ranging from 0, 50, 250 and 500 µM. The dose dependent reduction of colony spreading with 5HI was observed. All the images of petri plates were taken by Canon EOS DSLR, 700 camera. Data presented in the histograms are mean values ± the standard deviation (n=6).

Due to lack of mutagenesis tools presently, we conducted the physiological assays to evaluate the inhibition of gliding motility phenotype of PMSZPI in presence of 5-hydroxyindole (5 HI). When the soft LB agar medium (0.35%) was supplemented with different concentrations of 5 HI, the PMSZPI colonies exhibited dose dependent reduction in the colony spreading (Fig. 2D). These observations were similar to that of *Cellulophaga lytica* which showed inhibition of gliding motility in presence of the indole derivative like 5 HI (Chapelais-Baron *et al*., 2018). The binding of 5HI to T9SS, which is integral to gliding machinery, was suggested as a likely mechanism for causing inhibition of gliding motility in *lytica* (Chapelais-Baron *et al*., 2018). *Chryseobacterium* PMSZPI was observed to be a gliding Bacteroidete similar to *C. lytica* and *F. johnsoniae*. PMSZPI cells did not show any inhibition in their growth in presence of 5HI (Fig. S2) suggesting the latter’s non-toxicity towards the bacterial growth.

### Spreading colonies show remarkable cellular organization

Behaviour of PMSZPI cells in a spreading colony was visualized in detail using time lapse microscopy by inoculating 1μl of PMSZPI cells in the center of agar surface on a glass slide. For optimal spreading, 1/10 LB with 0.35% agar was used as the standard for our subsequent experiments. Following the inoculation of soft agar with PMSZPI cells and incubation for 2 h, time lapse imaging showed a rapid progression of the leading edges of the spreading colony (Fig. 3A). A closer examination of a leading edge migrating across the agar surface, revealed the organization of the cells in multiple layers-outermost layer appeared to be relatively transparent as compared to inner layers (i. e. towards the centre of the colony) (Fig. 3B). This might be due to thinly dispersed population of cells resulting in less cell density at the advancing front as compared to the inner layers which displayed relatively more closely packed rafts of cells.

**Figure 3:**
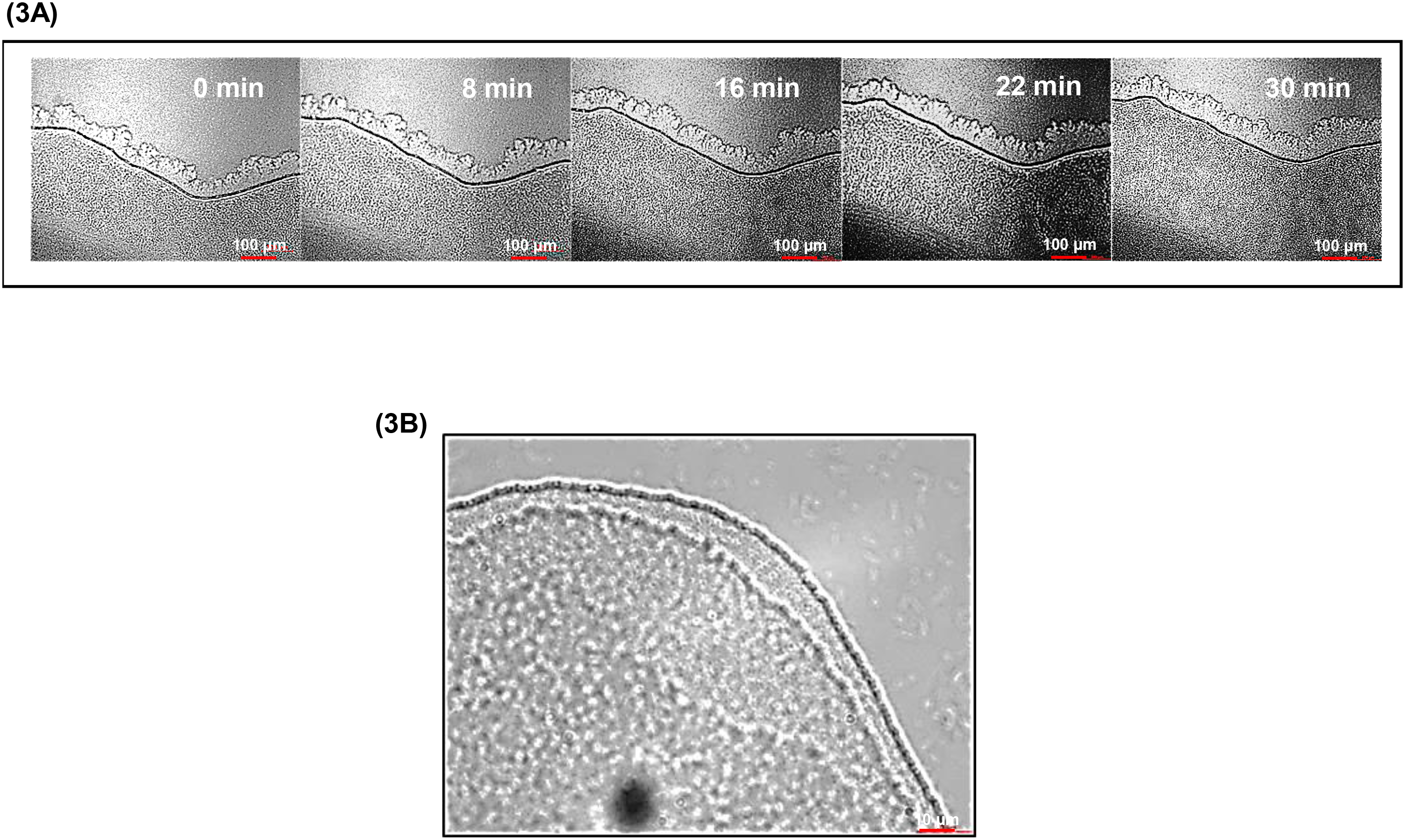
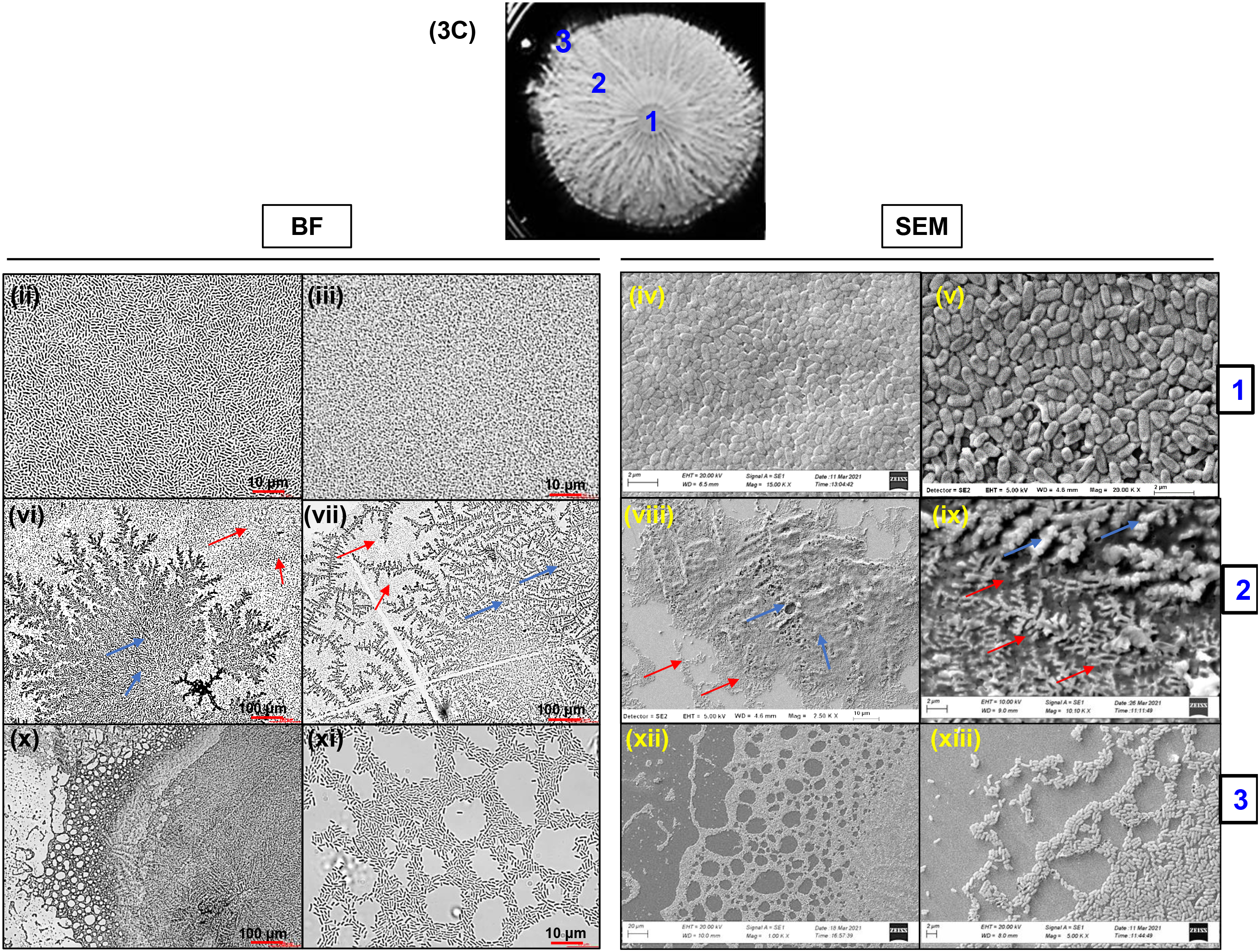
Structural characterization of spreading colonies. (A) Time lapse microscopy of development of colony edges. PMSZPI cells (1μl at cell density of ∼2×10^4^) was spotted on glass slide covered with a layer of LB (1/10) 0.35% agar, incubated for 2 h at 30°C and analysed *in situ*. The images of spreading edges of the growing colony were recorded with video camera attached to the microscope as described in Methods at regular intervals till 30 min. Progression of the spreading over time is visualized here. (B) Analysis of a leading edge. Higher magnification of a leading edge from A with bright field (BF) microscopy. The image displays the layered arrangement of the cells-the tip appearing transparent possibly due to less cell density followed by tightly packed cells towards interior. (C) Cellular organization in the spreading colony. Microscopic analysis of the cellular arrangement at different areas of spreading colony marked as 1, 2 and 3 was done. (i) Petri plate in Fig. 2B showing spreading colony of *Chryseobacterium* on LB (1/10) 0.35% agar. (ii) and (iii) are bright field micrographs from center of the colony depicting region of 1 whereas (iv) and (v) are the corresponding SEM images. This region shows dense packing of the cells. (vi) and (vii) presents bright field micrographs from region of 2 and (viii) and (ix) are the corresponding SEM images. This region shows the branched structures wherein the cells (red arrows) are interspersed within the extracellular matrix (blue arrows). (x) and (xi) are bright field micrographs from the edges of the colony from region of 3 whereas (xii) and (xiii) are the corresponding SEM images. The edges show the cells periodically arranged in hexagonal lattices.

We characterized the cell organization of the PMSZPI spreading colony grown for 24 h on soft agar (0.35%) to gain further insights into the structural arrangement of the cells by using bright field microscopy and scanning electron microscopy (Fig. 3C). The structural characterization was carried out at various regions within the spreading colony (Fig. 3Ci). The cells appeared to be densely packed and clustered forming multiple layers at the center of the colony (Fig.3Cii-v). The cell migration corresponded with the organized branched structures observed to be emanating radially from the center of the colony. Bright field microscopy of the branching region showed interesting ‘fern’ like branched structures beneath which the cells were found to be uniformly distributed (Fig. 3Cvi-vii). It seemed that the migration of the cells was associated with the packing and thickening of the branches by the multiplication of the cells (Fig. 3Ci). On further examination of the branching region with SEM, PMSZPI cells were found to be buried within a matrix of branched network which provided the impression of ‘fern’ like pattern suggesting the formation of biofilms (Fig. 3Cviii-ix). There lies a possibility that the PMSZPI cells were able to glide on the agar surface through the formation of extracellular matrix. Biofilm formation was observed in the spreading colonies of *F. johnsoniae* on 0.3% agar (Sato *et al*., 2021). Light microscopic images of the edges of the spreading colony showed clusters of cells periodically arranged in ‘honeycomb’ like patterns (Fig 3Cx-xi). A closer examination with SEM showed the cells organized into hexagonal lattices interconnected with each other correlating with the bright field microscopic imaging of the colony edges (Fig. 3Cxii-xiii). Such detailed image analysis provided a unique optical fingerprint of the spreading colonies of PMSZPI which is attributed to the bacterial ability to self-organize into various domains within a colony-closely packed at the centre and beneath the extracellular matrix in the branching region and as interconnected hexagonal lattices at the colony edges. Overall, our results present a multifaceted ordered structural organization of the cells in a spreading colony of the gliding PMSZPI.

### PMSZPI colonies show structural/iridescent coloration

Gliding motility has been strongly linked to structural coloration or iridescence in the gliding bacteria of Bacteroidetes phylum wherein it is suggested that gliding motility is required for the establishment of periodic structures within the iridescent colonies (Kientz *et al*., 2012a). The spreading colonies of PMSZPI (Fig. 2B) when visualized under trans-illumination with natural light exposure conditions displayed bright structural coloration (Fig. 4A). Iridescence could also be visualized in PMSZPI colonies by direct oblique illumination (data not shown). The coloration could have resulted from interaction of natural light with periodic cellular organization as visualized in PMSZPI colony (Fig. 3C) which has been commonly reported within Flavobacteria (Kientz *et al*., 2012b; Kientz *et al*., 2016; Johansen *et al*., 2018; Hamidjaja *et al*., 2020). The corresponding PMSZPI colonies on soft agar (0.35%) with medium concentrations ranging from 1/2 LB to 1/50 LB (Fig. 2B) exhibited higher levels of iridescence at lower nutrient conditions (Fig. 4A) possibly due to higher motility under nutrient deficient conditions (Fig. 2B). Such higher intensity iridescence profiles with low nutrient concentrations (Fig. 4A) could be visualized due to the ability of PMSZPI to self-organize into systematic lattice at the spreading edges (Fig. 3C x-xiii) or in distinct ordered layers in entirety within the colonies (Fig. 3Cii-ix) causing interference of the incident light. Highly motile *Flavobacterium* cells exhibiting iridescence organized into comprehensive periodic structures on low nutrient plates in contrast to those with reduced motility that could not show such organization (Johansen *et al*., 2018). Low iridescence coloration observed at higher LB (1/2) concentration could be attributed to low motility and low optical reflection possibly due suppression by yellow flexirubin pigment of PMSZPI. We further analyzed the development of structural colours under different culture conditions including agar concentrations and incubation time. Iridescent colors (corresponding to the colonies in Fig. 2C) were diminished at higher agar concentrations (0.5-1%) which seemingly reduced the motility and did not show any spreading edges (Fig. 4B). The colonies expanded with the increase in incubation period and such colonies exhibited strong coloration especially at the spreading edges (Fig. 4C). The peripheral corrugated edges of the expanded colonies showing structural coloration appeared to be layered with lower cell densities towards the extreme exterior (Fig. 4C).

**Figure 4:**
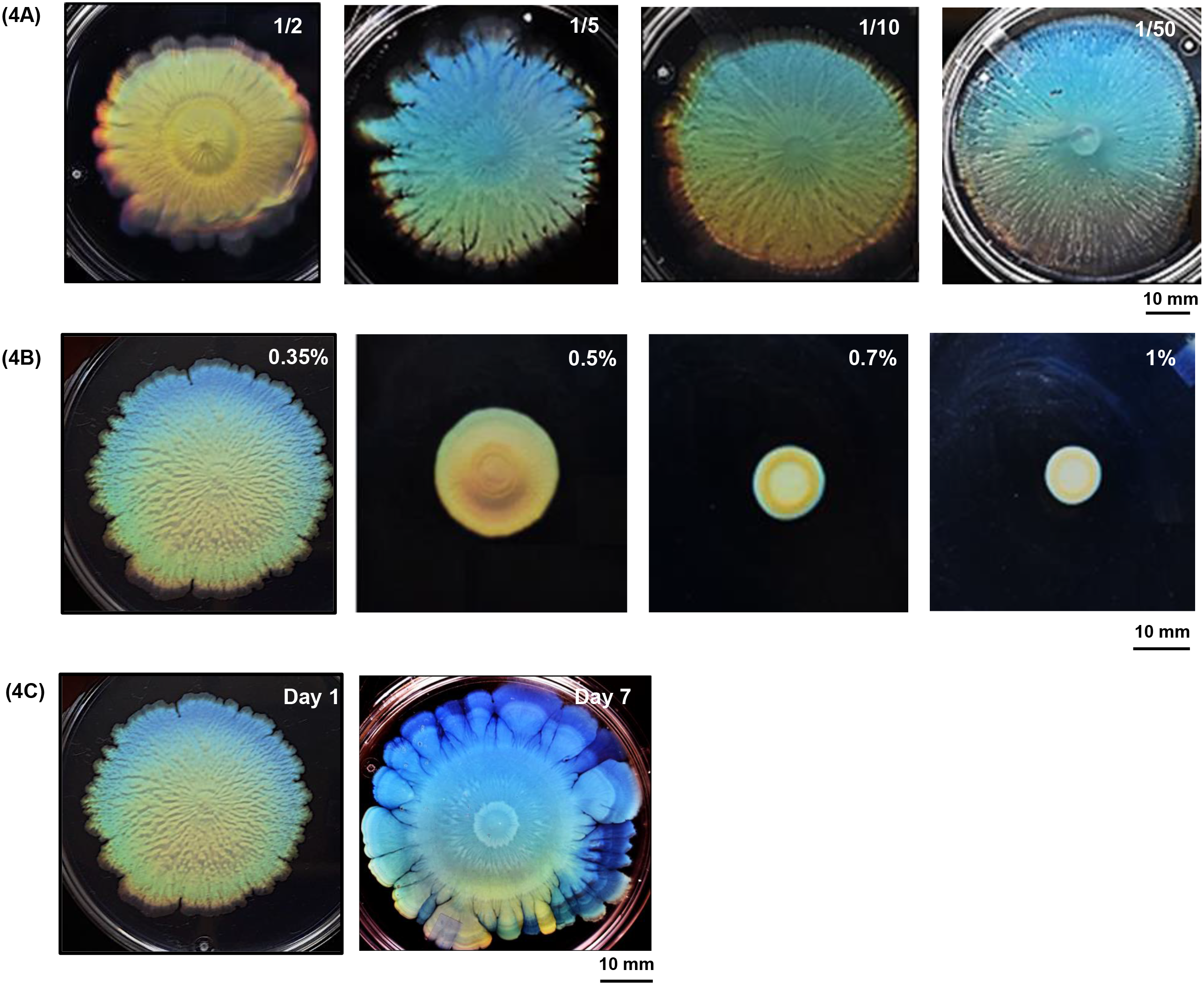
Structural/iridescent coloration in spreading colonies. Photographs of spreading colonies showing iridescent colors under transillumination in presence of (A) different LB concentrations (1/2, 1/5, 1/10 and 1/50) corresponding to plates shown in Fig. 2B. Scale corresponds to 10 mm. (B) different agar concentrations (0.35, 0.5, 0.7 and 1 %), corresponding to plates shown in Fig. 2C. (C) Iridescence of the spreading colony on 1/10 LB (0.35% agar) after 1 and 7 d of incubation. Iridescence was higher in the colonies that showed higher motility in lower LB and agar concentrations.

Generally, intense green iridescence was observed in marine Bacteroidetes and *F. johnsoniae* (Kientz *et al*., 2012a, b; Johansen *et al*., 2018). The nature of culture media played an important role in the colonial coloration(Kientz *et al*., 2012a, b). In our studies, we mostly observed the color gradation from blue, green, yellow and red with the spreading colonies with LB agar (Fig. 4). Bacterial iridescence is unknown in natural ecosystems. The structural coloration is proposed to attribute towards photoprotection or thermoregulation in marine Bacteroidetes members (Kientz *et al*., 2012a) or optimum cellular organization to degrade biological polymers (Johansen *et al*., 2018) or predation (Hamidjaja *et al*., 2020) in the soil bacterium, *F. johnsoniae.* Our work here provides the first evidence for iridescence in the Bacteroidetes gliding bacterium, *Chryseobacterium* PMSZPI isolated from uranium/metal enriched environment. Iridescence appears to be secondary consequence of gliding motility. It is suggested that the highly organized cell population with superior packing density in iridescent colonies of PMSZPI can be useful for degradation of biological polymers for nutrition or escaping from predation apparently conferring a survival advantage upon PMSZPI in such metal contaminated environment.

### PMSZPI adheres to glass surface and forms biofilm

Gliding motility has been implicated to be essential for bacterial attachment and colonization of plant surfaces in *Flavobacterium* spp. (Kolton *et al*., 2014). Moreover, components of type IX secretion system (T9SS) like GldK and SprT that were shown to be involved in the gliding motility and biofilm formation in the gliding Bacteroidetes bacterium like *Capnocytophaga ochracea* (Kita *et al*., 2016) were also harbored by PMSZPI. We therefore explored the ability of the gliding PMSZPI cells for attachment to the glass surface and biofilm formation. The cells spotted on glass slide following incubation for 5 min and three brief washes with the medium (Kita *et al*., 2016) when subjected to bright field microscopy and scanning electron microscopy revealed their firm attachment to the glass surface (Fig. 5). The cells following washes appeared to be typically organized in coordinated, regular clusters, lying side by side (Fig. 5B and C). In contrast, the cells spotted on the slide without washes appeared uniformly scattered on the glass surface (Fig. 5A). The cell attachment to glass surface was also evaluated quantitatively using Petroff-Hausser counting chamber to present consistent volume and concentration of cells. The average number of cells attached per field following washes (from 12 random fields, each field of 0.0025 mm^2^) was comparable to those found on glass surface without washes (Fig. 5D). In contrast, *E. coli* cells when evaluated in the similar way, did not show any attachment to glass surface when washed thrice with the medium (data not shown).

**Figure 5:**
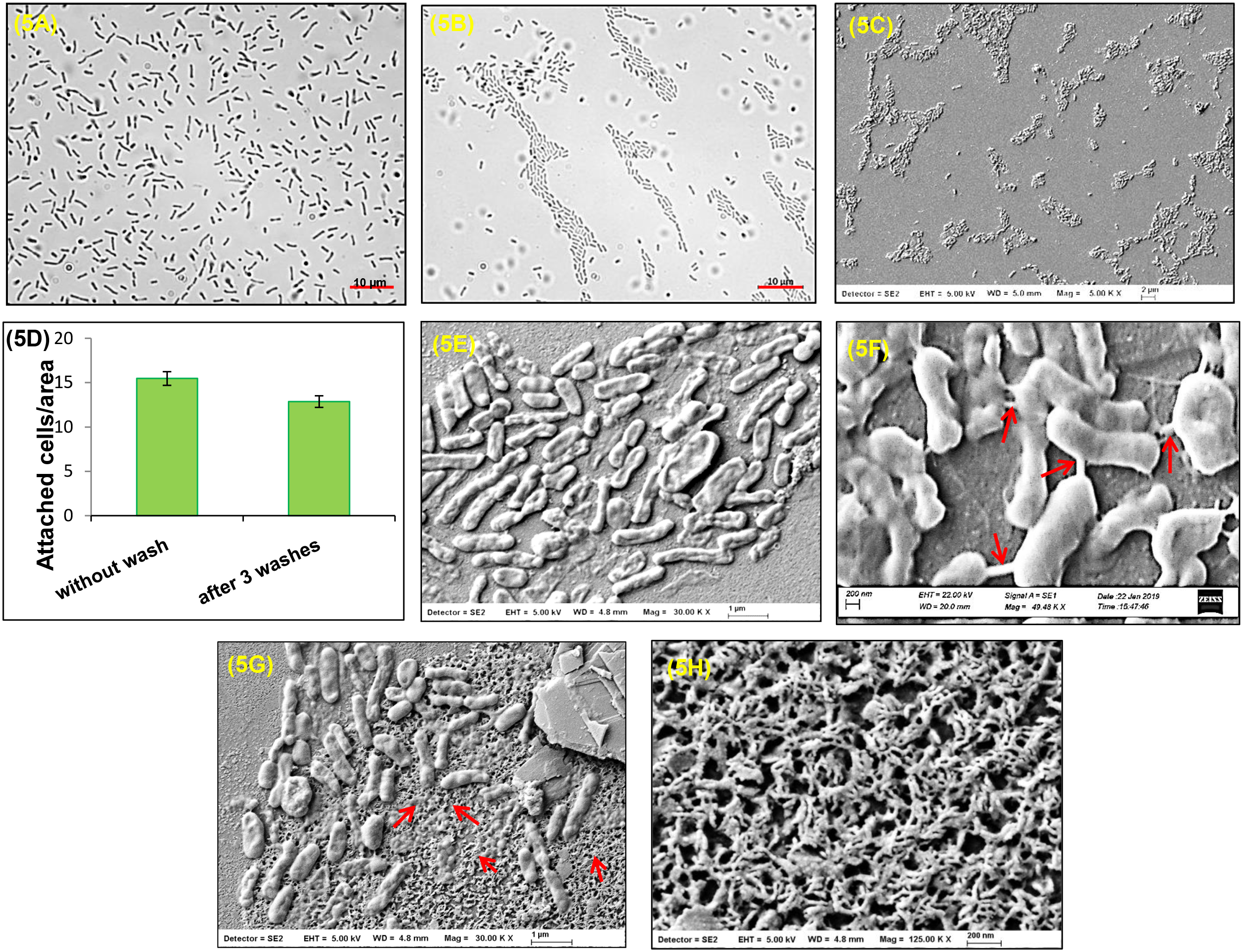
Cell attachment to glass surface and biofilm formation. (A) Microscopic analysis of cell attachment to glass surface. Bright field micrographs of cells spotted on glass slide, incubated for 2 min and visualized before washes and (B) after 3 washes with LB medium and (C) its corresponding SEM image. Uniform spreading of cells was observed before washes whereas the cells following washes showed aggregation lying side by side. (D) Quantification of cells attached to glass surface. Cells, added to Petroff-Hausser counting chamber and incubated for 2 min were washed thrice with LB medium. Shown here is the histogram depicting the number of cells attached to the glass surface (0.0025 mm2 region) before and after 3 washes were counted under microscope. Data presented are means ± SDs (n=12). (E) SEM analysis of biofilm architecture. The biofilms were cultured on glass slide for 5 days at 30°C followed by fixing, dehydration, gold coating and visualization by SEM. F. SEM image at higher magnification of the biofilm showing intercellular connections (arrows in red). G represents the formation of extracellular matrix (arrows in red) in the biofilm and H shows higher magnification of the fibrous extracellular matrix.

Attachment to glass surface is the initial step towards biofilm formation. As observed earlier, PMSZPI cells readily attached to glass surface (Fig. 5A-D). The ability of PMSZPI to form biofilms was evaluated by growing the cultures over glass slide for 120 h in 12-well plates. Scanning electron microscopy employed for analyzing the structures of the biofilms formed by the PMSZPI revealed biofilm formation with the cells closely packed together and interconnected with each other (Figs. 5E and F). In some areas of biofilm surface, fibrous extracellular matrix-like structures were also observed (Figs. 5G and H). Bacterial biofilms have been generally shown to be supported by extracellular polymeric substances (EPSs) which provides the mechanical stability to the biofilms (Flemming and Wingender, 2010). Biofilm formation assisted by extracellular matrix was also observed in the spreading colonies of PMSZPI (Fig. 3C, vi-ix). Overall, PMSZPI demonstrated the ability to attach to glass and form the biofilms similar to other Bacteroidetes members like *Capnocytophaga ochracea* (Kita *et al*., 2016) or *Flavobacterium* spp. (Kolton *et al*., 2014) harboring gliding motility genes.

### Uranium affects negatively on colony spreading and iridescence and promotes biofilm formation

We recently studied the genomic and functional diversities of PMSZPI which was isolated from uranium ore deposit and demonstrated its involvement in uranium bioremediation (Khare *et al*., 2020). In this study, we analyzed the effect of uranium on the gliding motility and consequently on iridescence of PMSZPI. The gliding motility responses of PMSZPI cells in the presence of uranium was evaluated on 1/10 LB medium supplemented with 0.35% agar. We chose 1/10 LB to avoid spontaneous precipitation of uranium. The motility and consequently the colony spreading of the cells were found to be dose dependently inhibited in presence of uranium as compared to control in absence of any metal (0-82% decrease for 0-200 µM U within 1 d) which was almost consistent over 7 days of incubation period (Figs. 6A and B). By the end of 7 days, although colony expansion was suppressed in presence of all tested concentrations of uranium, the colonies exhibited spreading edges representative of gliding bacteria (Fig. 6A). Iridescence was observed in the corresponding motility plates although the levels were lower (Fig. 6C) in comparison to control, U untreated plates (Fig. 4). Over 7 days, the iridescence was more concentrated on edges of the colonies (Fig. 6C). The higher cell densities as a result of growth over 7 days resulting in opacity of the colonies at the center could have resulted in the iridescence at the periphery with lesser cell densities. Time lapse microscopy of the advancing edge of the colonies in presence of uranium showed slow progression as compared to control (Fig. S3). Light microscopy and scanning electron microscopy analysis allowed for the investigation of cellular organization within the spreading colonies in response to uranium (25 µM for 24 h, Fig. 6Di). The close packing order of the cells was visualized at the center (Fig. 6Dii and iii) of the colony similar to that of control (Fig. 3C). We noticed ‘pores’ among the tightly packed cell population at the center by SEM (Fig. 6Dv) revealing multiple layers of the cells apparently contributing towards biofilm formation. It was reported that motile cells may enter into the mature biofilms, generating transient pores that enhance the nutrient flow in the matrix (Houry *et al*., 2012). However, the phenomenon of pore formation in our studies remains to be identified and is a subject of future studies. The spreading edges of colony showed ‘mesh’ like appearance (Fig. 6D iv) which on closer examination revealed the cell arranged periodically in lattice like structures (Fig. 6D vi and vii). Such packing may have given rise to angle-dependent optical response or iridescence in PMSZPI (Fig. 6C). Bacterial iridescence, otherwise unknown in natural environment, was conserved under conditions representing stressful marine ecosystems (Kientz *et al*., 2012a). As far as the authors are aware, bacterial iridescence in response to metals has not been investigated. In higher organisms such as feral pigeon, exposure to lead reduced the iridescent neck feather brightness possibly due to disruption in the production or arrangement of the microstructural feather elements, including melanosomes, needed for maximum colour expression (Chatelain *et al*., 2017). In our studies, uranium inhibited the motility of PMSZPI thereby reducing the levels of iridescence.

**Figure 6:**
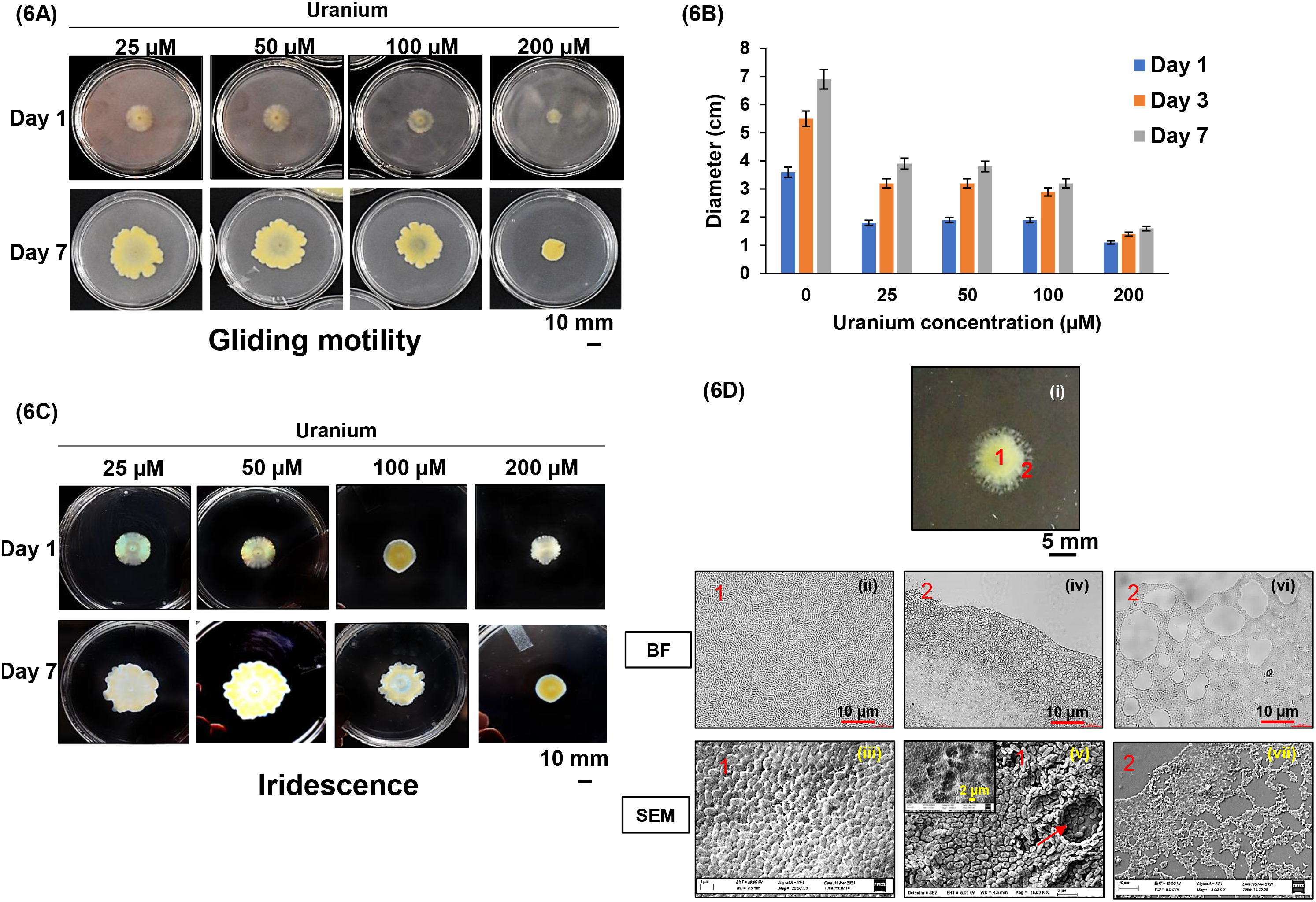
Influence of uranium on colony spreading and iridescence. (A) Colony spreading in presence of uranium. PMSZPI cells (10 µl at cell density of 2 x 10^5^ total cells) were spotted on LB (1/10) 0.35% agar supplemented with uranium (25-200 µM) and incubated at 30°C until 7 d. Shown here the images of colony spreading at different concentration of uranium on day 1 and day 7 and (B) shows the histogram depicting the colony diameters with progression of incubation time. Data presented here are mean values ± the standard deviation (n=6). There is significant decrease in the colony spreading in presence of increasing concentrations of uranium. (C) represents the iridescent coloration in plates corresponding to A. Iridescence decreases as the uranium concentration increases. (D) Colonial organization in presence of uranium. The cellular organization was visualized by BF and SEM at the centre marked as 1 and edges of the colony marked as 2. The dense packing of cells at the centre, showing the formation of pores (inset) or the periodic arrangement of cells in hexagonal lattices connecting to each other at the edges were similar to the control plates without uranium.

Motility in microbes is essential for escaping the toxic compounds in their immediate environment. It is suggested that if the cells detect any toxic compound, they can either avoid toxicity by initiating a motility process or adhere to a surface and form biofilms. We observed suppression of motility in presence of uranium. Therefore, we explored the process of biofilm formation by PMSZPI cells. The crystal violet staining method employed on biofilms grown on for 120 h in 12-well plates revealed significantly higher crystal violet associated biomass in presence of uranium (Fig. 7A). Almost ∼4 fold increase in biofilm formation was observed at 500 µM uranium as compared to uranium untreated control cells (Fig. 7 B). The structure of biofilms as examined by scanning electron microscopy in the presence of uranium over 120 h revealed denser packing of cells (Figs. 7C and D) interspersed within extracellular matrix (Fig. 7E) as compared to control (Fig. 5E). Reduced motility in presence of uranium could have resulted in denser biofilm in PMSZPI cells. Biofilm formation in response to uranium or any other heavy metal has not been explored in gliding bacteria yet. Furthermore, formation of biofilms could be an adaptation for PMSZPI cells for their survival in U enriched environment.

**Figure 7:**
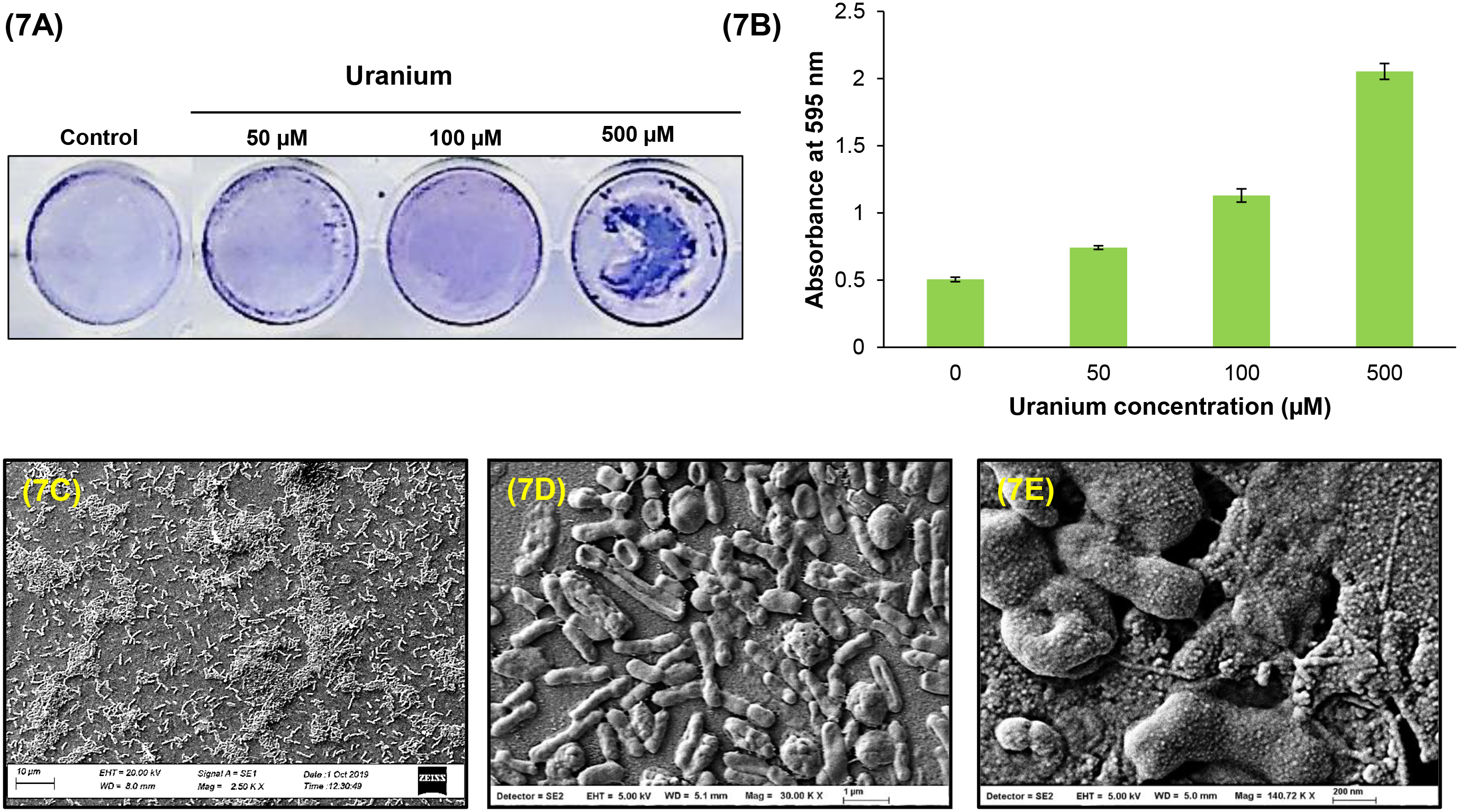
Biofilm formation in presence of uranium. (A) Crystal violet staining of biofilm. Cells were incubated on glass slides in absence and presence of uranium (50-500 µM) for 5 d at 30°C and imaged following crystal violet staining as described in methods. (B) Quantification of biofilm formation. The quantification of the biofilm produced was done by determining the OD_595_ following crystal violet staining. Increase in biofilm formation was observed with increase in uranium concentrations. Data presented here are mean values ± the standard deviation (n=6). (C), (D) and (E) are SEM images showing biofilm formation in presence of uranium at various magnifications.

## Conclusion

The ability of various bacteria to move over surfaces is a vital physiological characteristic that strongly supports their survival in their habitats. In this investigation, we demonstrate the distinctive features of the gliding motility of an environmental Bacteroidetes bacterium, *Chryseobacterium* sp. strain PMSZPI isolated from uranium enriched environment under different physiological conditions. The cellular organizational complexities that constitute the gliding process in PMSZPI resulting in spreading colonies were revealed in this study. The periodicity established within the gliding colonies gave rise to iridescence. It was observed that the presence of uranium caused inhibition to the gliding motility and iridescence and induced the formation of biofilms. Our studies discovered the key ecological processes like gliding motility and iridescence in this uranium tolerant soil bacterium that could be important for supporting its successful colonization and survival in otherwise hostile metal enriched ecosystem.

## Materials and methods

### Bacterial strain and culture conditions

*Chryseobacterium* sp. strain PMSZPI was isolated previously from sub-surface soil of the uranium ore deposit of Domiasiat in Meghalaya, India (Kumar *et al*., 2013). The PMSZPI preculture was initiated by streaking cells from a frozen 25% glycerol stock onto Luria-Bertani (LB) (Difco) agar plates and incubated overnight at 30°C. A single colony from the plate was inoculated into 10 ml of Luria-Bertani (LB) medium (Difco) in 25 ml borosilicate flask and incubated overnight under shaking (120 rpm) and aeration at 30°C. Such overnight grown cells were then inoculated into LB medium to initiate experiments. The ultrastructural analysis was performed by transmission electron microscopy and scanning electron microscopy as described earlier (Khare *et al*., 2020).

### Phylogeny and sequence analyses

Few representative strains of Phylum Bacteroidetes were used for constructing a maximum likelihood 16S rRNA tree. ClustalW (Thompson *et al*., 2003) was used for multiple sequence alignment of 16S rRNA gene sequences followed by generation of Maximum likelihood (ML) phylogenetic tree with 500 bootstrap replications using MEGA 7 v7.0.18 (Kumar *et al*., 2016). The orthologs to the gliding motility genes of *F. johnsoniae*, *gldB*, *gldD*, *gldH*, *gldJ*, *gldK*, *gldL*, *gldM*, *gldN*, *sprA*, *sprE*, and *sprT* were identified in the genomes of seven members of the phylum *Bacteroidetes* namely *Chryseobacterium sp.* PMSZPI*, Riemerella anatipestifer, Flavobacterium johnosoniae, Capnocytophaga orchracea, Cytophaga hutchinsonii, Prevotella melaninogenica,Cellulophaga lytica and Porphyromonas gingivalis* (Table S1) by BLAST analyses ( E values were set at 1e-5) and were confirmed as reciprocal best hits. Protein phylogenies (GldK, GldL, GldM, GldN) were evaluated using maximum likelihood (ML) and the reliability of individual tree was confirmed with 500 bootstrap replications.

### RNA isolation and Reverse transcriptase PCR (RT-PCR)

Overnight grown culture of PMSZPI cells in LB were used for the isolation of genomic DNA and RNA. Genomic DNA isolation was done by using the DNA isolation kit (BRIT, JONAKI, India). RNA isolation was done by using Illustra^TM^ RNAspin Mini kit (GE Healthcare Life Sciences, UK) as per the manufacturer’s instructions. RNA quality was checked by visualising 16S and 23S rRNA bands on 1% agarose gel. RNA concentrations (OD_260_) and purity (OD_260/280_) were determined by microplate reader (Biotek, Germany). DNA contamination were checked ahead of cDNA preparation. RNA was used to prepare cDNA by ReadyScript™ cDNA Synthesis Mix (Sigma) according to manufacturer’s instruction. RT PCR was employed to determine the transcriptional organization of *gldK, gldL, gldM and gldN*. PCR amplification by Taq polymerase (NEB) using primers for internal regions of KL, LM, MN and LN were done with genomic DNA, cDNA and RNA (negative control) as template.

### Colony spreading

PMSZPI cells were assessed for their movement over agar surfaces resulting in colony spreading. The cells were grown in LB broth at 30 °C with shaking (120 rpm) overnight. The cultures were harvested by centrifugation at 10,000 rpm for 3 min and were adjusted to an OD600nm∼1 with fresh LB. Aliquots of 10 μl (2 x 10^5^ total cells) from the resulting cell suspension were spotted onto the centre of agar medium in petri plates (9 cm in diameter) and incubated under various physiological conditions including incubation time (1-7 d), LB concentrations 1/ 2 (10 g l^-1^), 1/5 (4 g l^-1^), 1/10 (2 g l^-1^), 1/50 (0.4 g l^-1^)), agar concentrations 0.35% (3.5 g l^-1^), 0.5% (5 g l^-1^), 0.7% (7g l^-1^) and 1% (10 g l^-1^), and concentrations of 5-Hydroxyindole (5 HI) (0-500 µM) as indicated in the text. All plates were incubated at 30°C. Following requisite incubation, colony diameters were recorded and the petri plates were photographed by digital camera (Canon EOS DSLR, 700). For growth studies, exponential phase PMSZPI cells (OD600nm∼0.1) were added to sterile 2 ml LB medium amended with 50, 250 and 500 μM of 5-Hydroxyindole in polystyrene 12 well microplates. The growth was measured in terms of optical density at 600 nm using the Bio-Tek® Synergy^TM^ HT Multi-Detection Microplate Reader (Germany). Stock solution (100 mM) of 5-HI was prepared in methanol. Effect of uranium (U) on colony spreading was conducted on 1/10 LB (2g l^-1^) 0.35% agar in presence of different concentrations of uranyl carbonate (Acharya *et al*., 2009) ranging from 0-200 µM U.

### Time lapse microscopy

Cells of PMSZPI were grown overnight in LB medium and adjusted to OD600nm∼1with LB and 1 µl of suspension was spotted onto glass slides that had been earlier covered with a thin layer of LB (2g l^-1^) 0.35% agar medium. The cells were incubated at 30°C for 2 h and thereafter the edges of the colony were visualized by bright field microscopy (Carl Zeiss Axioscop 40 microscope with a charge-coupled device CCD Axiocam MRc Zeiss camera) and imaged at various time intervals mentioned in the text.

### Structural characterization of the spreading colony

The arrangement of the cells in the spreading colonies was characterized using bright field microscopy and scanning electron microscopy. The coverslips (5 mm dimeter) were placed on top of the colony at different locations. After 15 mins, the coverslip was picked up with forceps carrying the colonial impressions adhering to the coverslips. Subsequently the cells adhering to the coverslips were fixed with 2.5 % glutaraldehyde and were observed by bright field microscopy under oil immersion objectives (Carl Zeiss Axioscop 40 microscope with a charge-coupled device CCD Axiocam MRc Zeiss camera). For SEM, post fixation, the cells were serially dehydrated in 20, 30, 50, 70, 90 and 100% ethanol, sputter coated with gold and observed using Zeiss Evo 18 SEM (UK) as well as field-emission scanning electron microscopy (FE-SEM) (Carl Zeiss Auriga, Germany).

### Iridescence profile

Iridescence of bacterial colonies exhibiting spreading was observed under transillumination. Initially, the petri plates with the colonies were tilted to allow the light to shine through and were visually examined for structural coloration. Subsequently, the colonies exhibiting iridescence were photographed from an angle of 45° above the petri plates with the light source (natural light) directly behind it (Kientz *et al*., 2012a).

### Bacterial attachment to the glass surface

The stationary phase culture of PMSZPI was adjusted to an OD_600nm_ of 1 with fresh LB broth and 10 µl of the cell suspension was added to glass slide. After 2 min of incubation, three brief washes with LB (200 µl of medium) were given to the cells to remove the unattached cells. Cells which remained attached to the slides following washes were visualized by bright field microscopy under oil immersion objectives (Carl Zeiss Axioscop 40 microscope with a charge-coupled device CCD Axiocam MRc Zeiss camera) and scanning electron microscopy (Zeiss Evo 18 SEM, UK).

Quantification of the attached cells to the glass surface was done using a Petroff-Hausser counting chamber as described earlier (Nelson *et al*., 2007) with some modifications. Overnight grown cells of PMSZPI were freshly inoculated in LB medium and incubated at 30°C to attain the OD_600nm_ of 0.3. Aliquot of 2.5µl was added to Petroff-Hausser counting chamber and incubated for 2 min at room temperature. After 2 min of incubation, unattached cells were removed by three brief washes with LB (200 µl of medium) and the remaining cells were covered with a coverslip. The number of cells attached to 12 randomly selected 0.0025 mm^2^ regions of the chamber was counted using bright field microscopy.

### Biofilm formation

Crystal violet assays for quantification of biofilm in absence and presence of uranium were done in a polystyrene 12 well microtiter plates in three wells for each condition. The stationary phase cells were adjusted to OD600nm∼0.5 with fresh LB and added to the wells having 1/10 LB medium (2 ml/well) without or with uranium (0-500 µM uranyl carbonate). After incubation for 5 days at 30° C, the wells were washed with distilled water and subsequently stained with 0.1% crystal violet for 10 min at room temperature. Following staining, the wells were washed twice with distilled water and the plate was allowed to air-dry. The biomass associated with crystal violet was extracted with acetic acid (30%) and transferred to a new plate. The absorbance was measured at 595 nm with Bio-Tek® Synergy^TM^ HT Multi-Detection Microplate Reader (Germany).

Scanning Electron Microscopy (SEM) was used to study the biofilm structure in absence and presence of uranium (100 µM). Glass slides were placed in the wells of polystyrene 6-well microtiter plates with 4ml/well of 1/10 LB medium. The stationary phase cells were adjusted to OD600nm∼0.5 with fresh LB and were added to the wells and incubated at 30° C for 5 d under static conditions. Thereafter, the slides were washed with saline solution and fixed with 2.5 % glutaraldehyde. Post fixation, the cells were serially dehydrated in 20, 30, 50, 70, 90 and 100% ethanol. The slides were gold coated and visualized using Zeiss Evo 18 SEM and field-emission SEM (FE-SEM)) (Carl Zeiss Auriga, Germany).

## Acknowledgements

The authors thank Dr. H.S. Misra, Head, Molecular Biology Division, BARC for his constant support and encouragement during the course of this study. This work was supported by Bhabha Atomic Research Centre, Department of Atomic Energy, Government of India.

