## Supplemental Table 1 for "Gliding motility of a uranium tolerant *Bacteroidetes* bacterium *Chryseobacterium* sp. strain PMSZPI: Insights into the architecture of spreading colonies"

**Table S1A: List of strains and NCBI accession numbers of genomes analysed for gliding motility related proteins**

| Strain | Genome accession number |
|---|---|
| *Flavobacterium johnsoniae* DSM 2064 | FRAO00000000 |
| *Chryseobacterium sp*. PMSZPI | PIZV01000000 |
| *Capnocytophaga ochracea* DSM 7271 | CP001632 |
| *Cellulophaga lytica* DSM 7489 | CP002534 |
| *Riemerella anatipestifer* ATCC 11845 DSM 15868 | CP003388 |
| *Cytophaga hutchinsonii* ATCC 33406 | FPJX00000000 |
| *Porphyromonas gingivalis* ATCC 33277 | AP009380 |
| *Prevotella melaninogenica* ATCC 25845 | CP002122, CP002123 |

**Table S1B: Sizes of *gldK, gldL, gldM, gldN* in *Chryseobacterium sp.* PMSZPI and other Bacteroidetes members**

| Nucleotide length (bp) | | | | | | | | |
|---|---|---|---|---|---|---|---|---|
| **Gene name** | ***Chryseobacterium* PMSZPI** | ***Flavobacterium*** | ***Capnocytophaga*** | ***Cellulophaga*** | ***Riemerella*** | ***Cytophaga*** | ***Porphyromonas*** | ***Prevotella*** |
| *gld K* | 1422 | 1395 | 1377 | 1341 | 1413 | 1035 | 1476 | 1482 |
| *gld L* | 693 | 648 | 666 | 654 | 687 | 786 | 930 | 798 |
| *gld M* | 1590 | 1542 | 1581 | 1560 | 1560 | 1596 | 1551 | 1590 |
| *gld N* | 939 | 990 | 900 | 903 | 915 | 885 | 1080 | 1044 |
