## Supplemental Table 2 for "Gliding motility of a uranium tolerant *Bacteroidetes* bacterium *Chryseobacterium* sp. strain PMSZPI: Insights into the architecture of spreading colonies"

**Table S2: Primer sequences used in the study.**

| Primers | Sequence (5’-3’) |
|---|---|
| Fwd KL | TGGCAAACTTTAAGCCTAAGCGA |
| Rev KL | AGCATGTTTATCTAATAATTCAGG |
| Fwd LM | CTGTTGACGTATCTGCTTCTA |
| Rev LM | TAGTAATAAAGCCAATGCATCA |
| Fwd MN | TTGTGAAACCTACAACAGGAACTA |
| Rev MN | TTAAGATATCGATAGCAGCAT |
| Fwd LN | CTGTTGACGTATCTGCTTCTA |
| Rev LN | TTAAGATATCGATAGCAGCAT |
