## Supplemental Figure 1 for "Gliding motility of a uranium tolerant *Bacteroidetes* bacterium *Chryseobacterium* sp. strain PMSZPI: Insights into the architecture of spreading colonies"

**Supplementary Figure S1: Phylogenetic relationship of sequences of gliding proteins.** The sequences for gliding proteins for the Bacteroidetes strains were obtained from NCBI and phylogenetic trees were constructed by maximum likelihood (ML) method using MEGA7 package with 500 bootstrap replications. The trees correspond to A. GldK, B. GldL, C. GldM and D. GldN. The proteins are shown by their accession numbers and the names of the organisms.

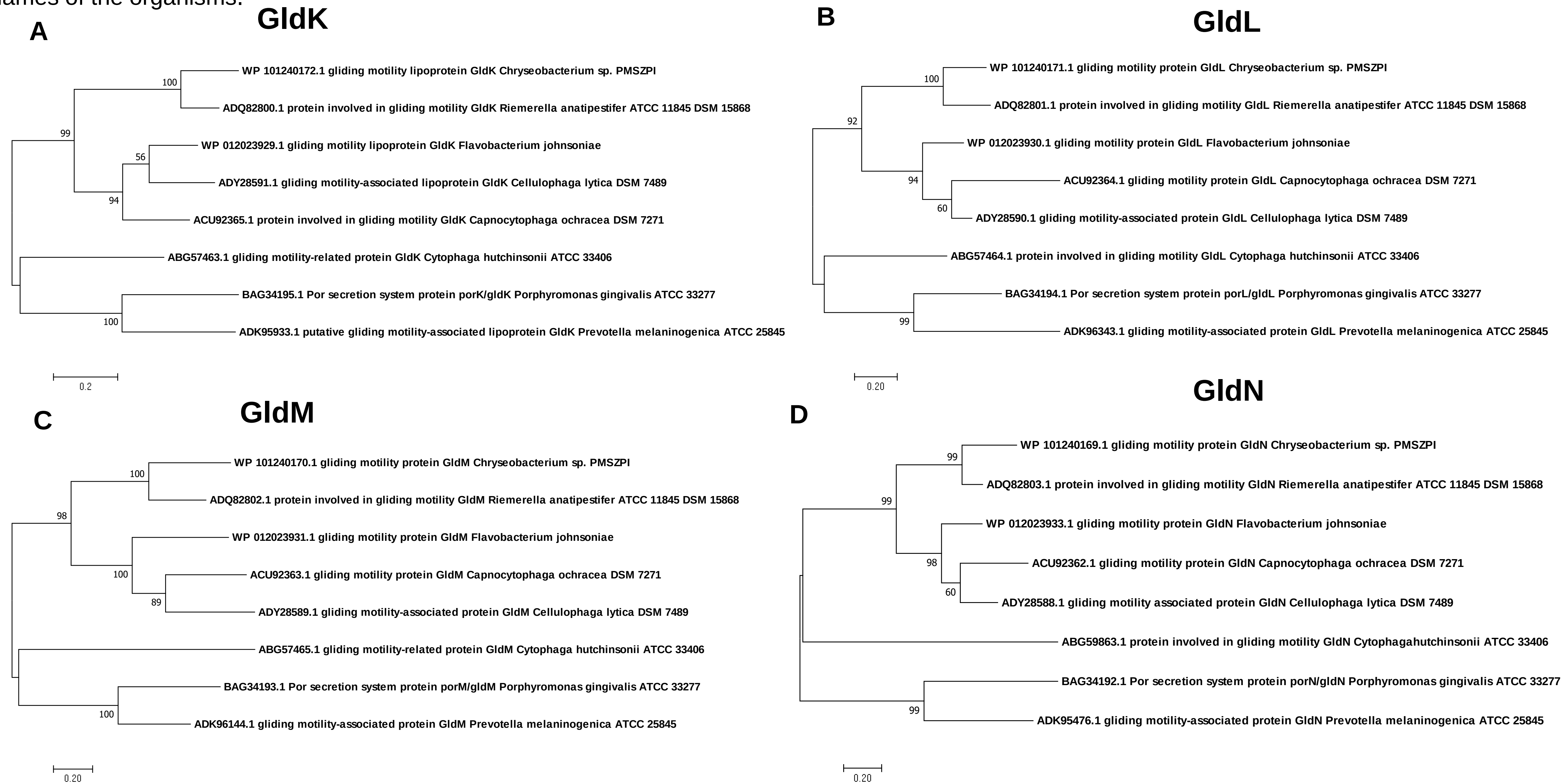
