## Supplemental figure 2 for "Gliding motility of a uranium tolerant *Bacteroidetes* bacterium *Chryseobacterium* sp. strain PMSZPI: Insights into the architecture of spreading colonies"

**Supplementary Figure S2: Effect of 5- hydroxyindole on growth of *Chryseobacterium* PMSZPI.** Growth studies were done for PMSZPI in absence or the presence of 50, 250 and 500 µM of 5- hydroxyindole at 30$^{o}$C .

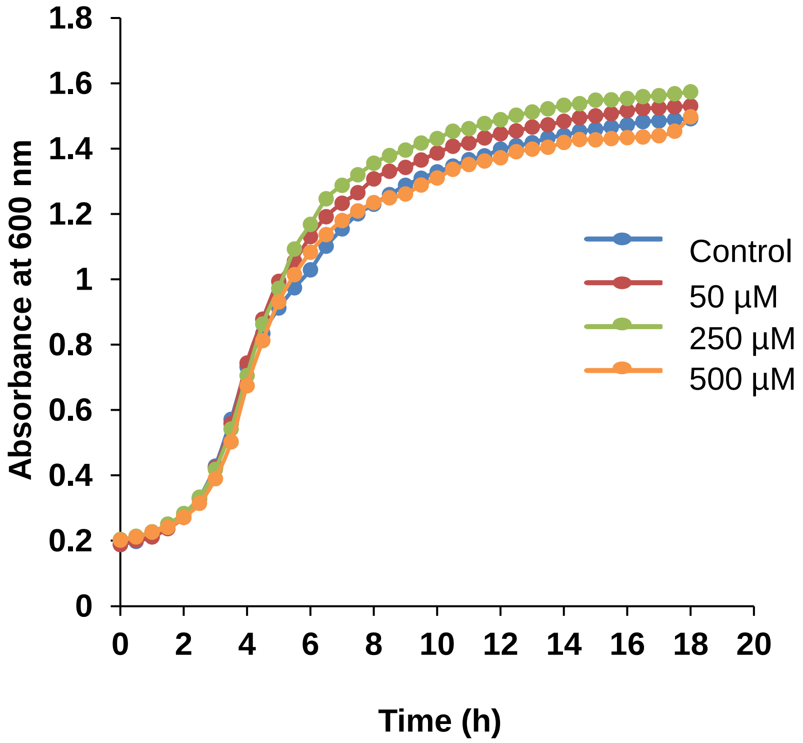
