## Supplemental figure 3 for "Gliding motility of a uranium tolerant *Bacteroidetes* bacterium *Chryseobacterium* sp. strain PMSZPI: Insights into the architecture of spreading colonies"

**Supplementary Figure S3: Time lapse microscopy of growing edge of PMSZPI colony in presence of uranium.** Growing edge of uranium exposed PMSZPI colony on 1/10 LB 0.35% agar was imaged at various time intervals mentioned in the text. Scale bar corresponds to 100 µm. Almost negligible movement of the edge(red arrow) was observed during 30 min.

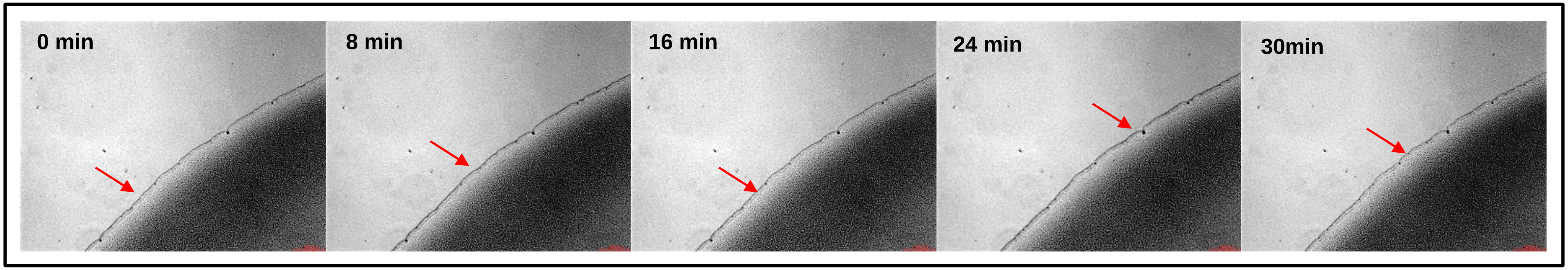
